# Rapamycin and caspofungin show synergistic antifungal effects in caspofungin-susceptible and caspofungin-resistant *Candida* strains *in vitro*

**DOI:** 10.1101/2023.10.05.560425

**Authors:** Maxime Lefranc, Isabelle Accoceberry, Nicolas Biteau, Thierry Noël

## Abstract

**Objectives:** Caspofungin is an echinocandin antifungal agent that inhibits synthesis of glucan required for the fungal cell wall. Resistance is mediated by mutation of Fks1 glucan synthase, among which S645P is the most common resistance-associated polymorphism. Rapamycin is a macrolide that inhibits the mechanistic target of rapamycin (mTOR) protein kinase activity. This study investigated the interaction between rapamycin and caspofungin in inhibiting the growth of wild type *Candida albicans* and Fks1 S645P mutant clinical isolate and wild type *Candida lusitaniae* and genetically engineered isogenic strain with Fks1 S645P mutation at equivalent position.

**Methods:** Interactions between caspofungin and rapamycin were evaluated using the microdilution checkerboard method in liquid medium. The results were analysed using the fractional inhibitory concentration (FIC) index and the response surface (RS) analysis according to the Bliss model.

**Results:** Synergy between rapamycin and caspofungin was shown for *C. albicans* and *C. lusitaniae* strains by RS analysis of the checkerboard tests. Synergy was observed in strains sensitive and resistant to caspofungin. Weak subinhibitory concentrations of rapamycin were sufficient to restore caspofungin susceptibility.

**Conclusions:** We report here for the first time synergy between caspofungin and rapamycin in *Candida* species. Synergy was shown for strains susceptible and resistant to caspofungin. This study highlights the role of the TOR pathway in sensing antifungal-mediated cell wall stress and in modulating the cellular response to echinocandins in *Candida* yeasts.

## Introduction

Invasive fungal infections are a major cause of global morbidity and mortality, accounting for nearly 1.4 million deaths every year (1). Echinocandins have become the first-line therapy for invasive candidiasis in most patients because of their excellent safety profile and good *in vitro* fungicidal activity against *Candida* species, including azole-resistant clinical isolates (2). They act as noncompetitive inhibitors of the fungal β(1,3)-glucan synthase, and thus disturb fungal cell wall synthesis (3). In *Candida albicans*, the main mechanism of echinocandin resistance involves nonsynonymous mutations in two hotspot regions of the *FKS1* gene, HS1 and HS2, which decrease the binding affinity of glucan synthase to echinocandins (4). Substitutions involving Ser-645 (S645P/F/Y) are the most common polymorphisms and are responsible for the most pronounced elevations in minimum inhibitory concentration (MIC) (5,6).

Tolerance and resistance to echinocandins have also been shown to involve regulatory pathways dependent on the Hsp90 chaperone and its client protein, calcineurin, in *C. albicans* (7). A previous study on *Candida lusitaniae* reported that the calcineurin inhibitor tacrolimus increased susceptibility to caspofungin of strains that were susceptible and resistant to echinocandins, notably in a strain harbouring an Fks1 mutation at the equivalent S645 position (8). The cellular receptor of tacrolimus is Fpr1p, a peptidyl-prolyl isomerase. Fpr1p also binds rapamycin, a natural product of *Streptomyces hygroscopicus*, which was first described as an antifungal agent capable of inhibiting *C. albicans* growth (9) but was later shown to have strong immunosuppressive activity (10). The cellular target of rapamycin is target of rapamycin complex 1 (TORC1), which is involved in the regulation of many cellular processes, including protein translation, autophagy, and stress responses (11,12).

To the best of our knowledge, few studies have been conducted on the impact of rapamycin activity on the sensitivity and resistance of *Candida* species to other antifungal agents (13). To gain insights into the possible interaction between the Fpr1–rapamycin complex and echinocandins, we evaluated the *in vitro* interaction between rapamycin and caspofungin against *C. albicans* wild type strain and Fks1 S645P mutant clinical isolate, and *C. lusitaniae* wild type strain and a genetically engineered isogenic strain with Fks1 S645P mutation at the equivalent position.

## Materials and methods

### *Candida* strains

The different strains used in this study are listed in Table 1. *C. albicans* ATCC 90029 and *C. lusitaniae* CBS 6936 (ATCC 38553) were used as susceptible reference strains. The echinocandin-resistant *C. albicans* strain was a clinical isolate obtained from the Laboratory of Mycology of Bordeaux University Hospital and identified by matrix-assisted laser desorption ionisation-time of flight (MALDI-TOF) mass spectrometry (MS) (Microflex LT system; Bruker Daltonics, Billerica, MA, USA). The nucleotide sequence of the entire *FKS1* gene was determined, and comparison with the sequence of the same gene of *C. albicans* ATCC 90029 revealed a heterozygous mutation T1933C resulting in substitution of a serine by a proline (S645P) in the HS1 region (amino acids 641–649) of glucan synthase (GenBank accession no. D88815). Then the same mutation was introduced at the equivalent position in the *FKS1* allele (T1912C substitution) in a genetically engineered strain of *C. lusitaniae* derived from the wild-type strain CBS 6936, as described previously (14).

**Table 1.**
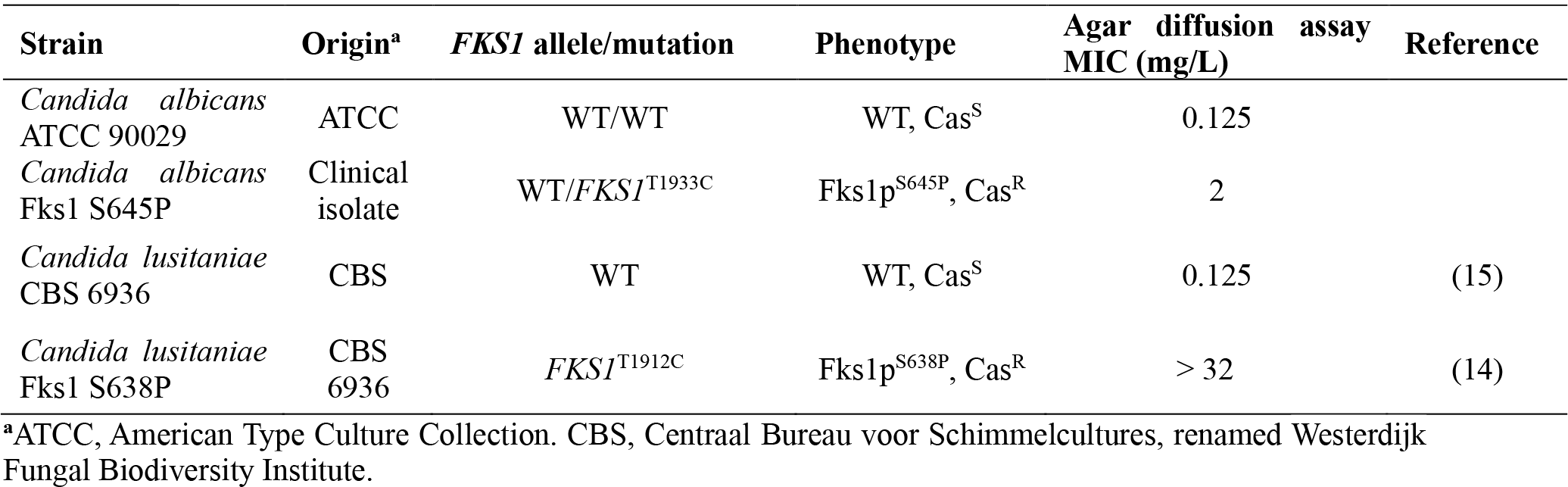
Name, genotype, phenotype, and caspofungin susceptibility of the strains used in this study.

### *In vitro* susceptibility testing

Stock solutions of 10 mg/mL caspofungin (Euromedex, Souffelweyersheim, France) and 10 mg/mL rapamycin (Euromedex) were prepared in dimethylsulfoxide (DMSO) and kept at – 20°C until use.

The checkerboard method used for drug combination studies was based on microdilution CLSI standards (16) in RPMI 1640, pH 7.0, buffered with 0.165 M 3-(*N*-morpholino)propanesulfonic acid (MOPS).

### MIC determination and interpretation

Caspofungin at concentrations ranging from 0.0625 to 4 mg/L was combined with rapamycin at concentrations ranging from 0.001 to 1 or 0.01 to 10 mg/L depending on the susceptibility of the strains. Yeast suspensions were diluted with RPMI 1640 at a final cell density of 1 × 10^3^ cells/mL in 96-well plates and incubated for 48 h at 35°C; growth was measured with a microplate reader at 450 nm. All experiments were performed in triplicate in independent assays, and growth variation did not exceed 10%. The MIC was defined at 90% growth inhibition for both drugs tested alone and in combination. High off-scale MICs were converted into the next twofold highest concentration. Two different methods were used to analyse the drug interactions: one based on the Loewe additivity model (calculation of the fractional inhibitory concentration index, FICI) and another based on the Bliss independence model (response surface modelling).

The FICI was calculated as follows: FICI = (MIC_combination_/MIC_alone_)_caspofungin_ + (MIC_combination_/MIC_alone_)_rapamycin_. The FICI data were interpreted as: synergy, FICI ≤ 0.5; indifferent, FICI > 0.5–4; and antagonism, FICI > 4.0 (17).

For response surface analyses, the experimental data generated were expressed for each well as a percentage of growth in the presence of drugs compared to the growth control in drug-free medium and then transformed into a dose–response curve for each drug alone. We used the Bliss independence model, which is based on the hypothesis that drugs act independently from each other. Using this model, a theoretical response surface of the combination corresponding to an indifferent interaction was calculated using the dose-response curves of both drugs. To calculate the synergy distribution, the modelled response surface was compared to experimental data. The effect of the drugs in combination was defined as synergistic or antagonistic if the observed effect lay below or above the predicted indifferent dose–response surface, respectively. All calculations were performed with Combenefit software (18). *In vitro* caspofungin susceptibility was also determined using Etest (bioMérieux, Marcy-l’Étoile, France) in accordance with the manufacturer’s instructions (Table 1).

## Results

Table 2 shows the results regarding the *in vitro* interaction between caspofungin and rapamycin against strains of *C. albicans* and *C. lusitaniae* susceptible and resistant to caspofungin. Caspofungin MICs and rapamycin MICs for each strain were within ± 2log_2_ dilutions in all experiments.

**Table 2.** *In vitro* interaction between caspofungin and rapamycin against strains of *C. albicans* and *C. lusitaniae* susceptible and resistant to caspofungin.

| Strains | Checkerboard MICs (mg/L) |  |  | FICI<sup>a</sup> |  | Response surface analysis |  |
| --- | --- | --- | --- | --- | --- | --- | --- |
|  | Caspofungin alone | Rapamycin alone | Caspofungin and rapamycin in combination | Value | INT | SUM-SYN-ANT | INT |
| *C. albicans* ATCC 90029 | 0.25 | 0.5 | 0.125/0.002 | 0.504 | IND | 49.77 | SYN |
| *C. albicans* Fks1 S645P | 2 | 0.5 | 0.125/0.25 | 0.765 | IND | 36.38 | SYN |
| *C. lusitaniae* CBS 6936 | 1 | 10 | 0.064/0.31 | 0.249 | SYN | 75.36 | SYN |
| *C. lusitaniae* Fks1 S638P | 8 | 10 | 1/0.31 | 0.27 | SYN | 69.53 | SYN |

<sup>a</sup>FICI, fractional inhibitory concentration index; INT, interpretation; SYN, synergistic; IND, indifference

The *C. albicans* and *C. lusitaniae* reference strains exhibited caspofungin MICs of 0.25 and 1 mg/L and rapamycin MICs of 0.5 and 10 mg/L, respectively. As expected, strains bearing the S645P mutation in Fks1 had higher caspofungin MIC values: 2 mg/L for *C. albicans* and 8 mg/L for *C. lusitaniae* (8 mg/L being the high-off scale MIC). The MICs of rapamycin for the *FKS1* mutant strains were identical to those observed in the corresponding reference strains (Table 2), i.e., 0.5 mg/L for *C. albicans* Fks1 S645P and 10 mg/L for *C. lusitaniae* Fks1 S638P. Analysis of the results of the checkerboard tests with the response surface approach based on the Bliss model revealed synergy between caspofungin and rapamycin for all strains. To visualise the results, the synergy levels were mapped onto the experimental combination dose– response surface. To summarise the synergy distribution, the SUM-SYN-ANT metric was used, which represents the sum of synergy and antagonism observed in the concentration range. As described elsewhere (19), for interpreting the SUM-SYN-ANT metric, a control plate with combinations of different concentrations of caspofungin alone was used for each reference strain of *Candida*. For *C. albicans* ATCC90029, the SUM-SYN-ANT of the control plate was 13.41%. Synergy and antagonism between caspofungin and rapamycin were assumed when the SUM-SYN-ANT was > 13.41% and < –13.41%, respectively. No drug interaction was considered for values between –13.41% and 13.41%. For *C. lusitaniae* CBS 6936, this metric was 25.11%.

Using caspofungin and rapamycin in combination, the SUM-SYN-ANT metric for *C. albicans* reference strain and *C. albicans* Fks1 S645P were 49.77% and 36.38%, respectively, indicating synergy between the two agents. For *C. albicans* Fks1 S645P, the presence of rapamycin at a concentration of 0.25 mg/L (i.e., half the MIC) reduced the MIC of caspofungin by a factor of 16 from 2 to 0.125 mg/L, thus restoring the sensitivity of the *C. albicans FKS1* mutant to caspofungin (Figure 1). The restoration of sensitivity to caspofungin in the presence of a low dose of rapamycin was confirmed in *C. lusitaniae* Fks1 S638P, in which a dose of 0.31 mg/L rapamycin (32 times less than the MIC) resulted in a decrease in caspofungin MIC from 8 to 1 mg/L (8 mg/L being the high-off scale MIC) (Figure 2).

**Figure 1.**
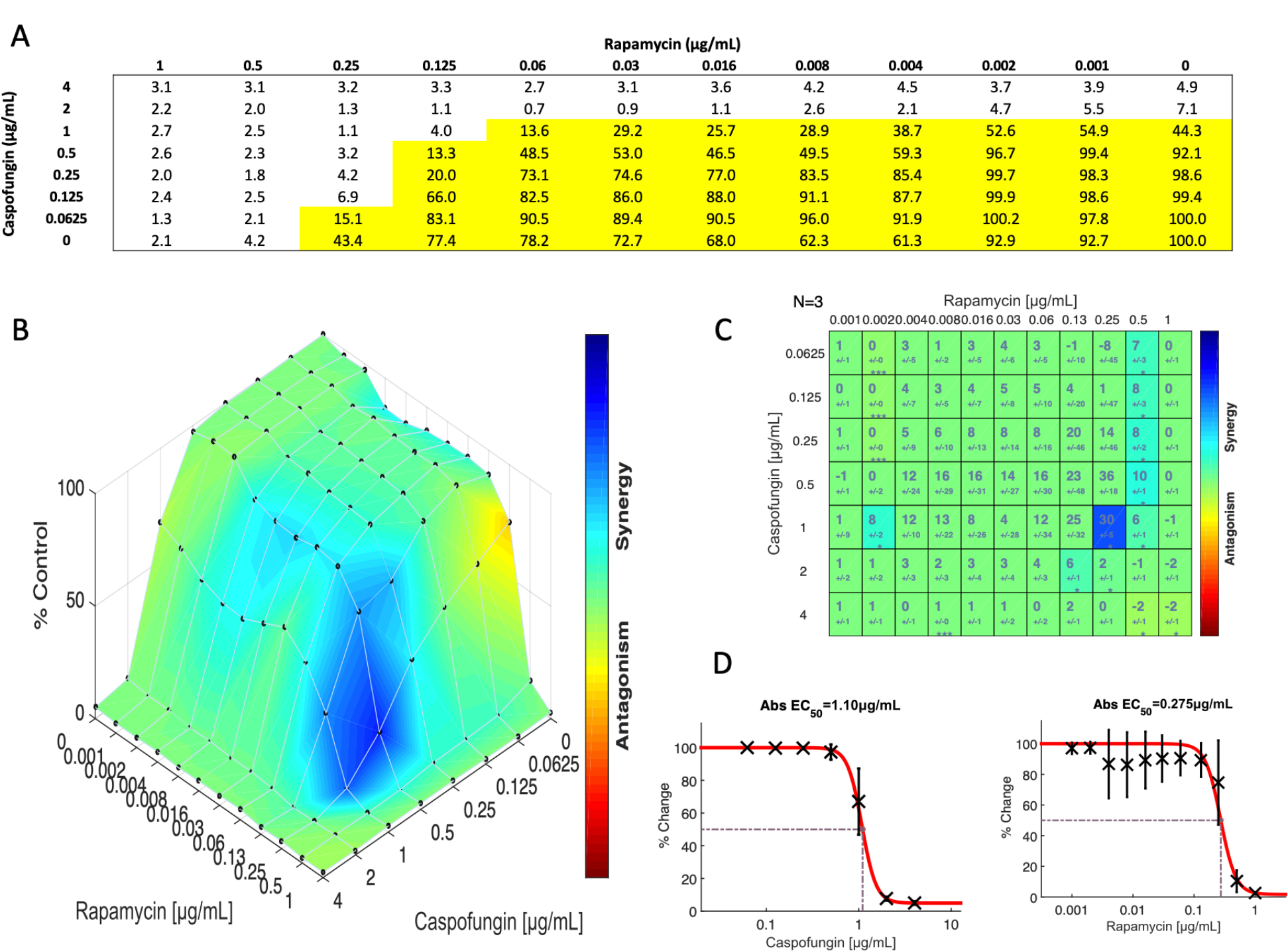
Checkerboard test using caspofungin and rapamycin with the *C. albicans* Fks1 S645P strain. (A) Percentage of yeast growth compared to the growth in drug-free medium on 96-well microplates. (B) Response surface analysis showing the mapping of the synergy levels based on the Bliss model. (C) Matrix of the synergy distribution derived from the combination dose–response and from the reference dose–response. (D) Dose– response curve of each drug alone. Panels B, C, and D were generated with Combenefit, version 2.021.

**Figure 2.**
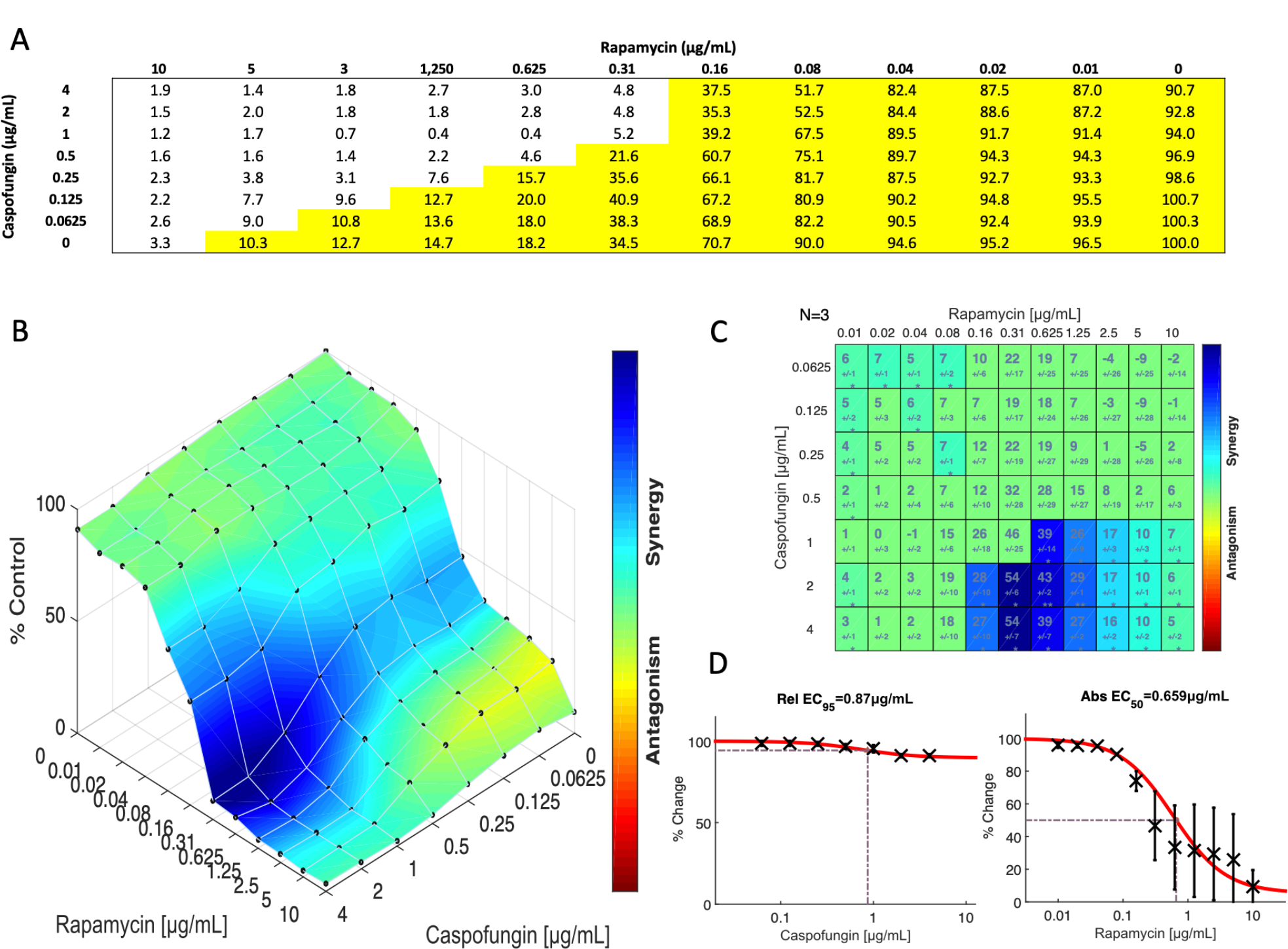
Checkerboard test using caspofungin and rapamycin with the *C. lusitaniae* Fks1 S638P strain. (A) Percentage of yeast growth compared to the growth in drug-free medium on 96-well microplates. (B) Response surface analysis showing the mapping of the synergy levels based on the Bliss model. (C) Matrix of the synergy distribution derived from the combination dose–response and from the reference dose–response. (D) Dose– response curve of each drug alone. Panels B, C, and D were generated with Combenefit, version 2.021.

This observation was confirmed by response surface analysis showing high SUM-SYN-ANT values of 75.36% and 69.53% for *C. lusitaniae* CBS 6936 and *C. lusitaniae* Fks1 S638P, respectively, reflecting a strong synergistic effect between caspofungin and rapamycin for both strains.

Analysis of the checkerboard results via FICI revealed synergy between caspofungin and rapamycin only for *C. lusitaniae* CBS 6936 and *C. lusitaniae* Fks1 S638P, with an FICI of 0.249 and 0.27, respectively. For *C. albicans* strains, even if the FICI values were low, ranging from 0.504 to 0.765, the interaction was considered as indifferent (Table 2). These results suggest that the FICI method is not sufficiently discriminating to detect synergistic effects when the MICs for both drugs are low.

## Discussion

The interactions between rapamycin and antifungals have been little studied, probably because rapamycin, first described for its antifungal properties (9), has also been shown to be an immunosuppressant (10) thus making its use for the treatment of opportunistic fungal infections inconceivable.

However, other immunosuppressants show synergistic activity with echinocandins or azole antifungals, such as ciclosporin or tacrolimus, which inhibit calcineurin (20–22). Tacrolimus binds to the Fpr1 protein, a peptidyl-prolyl isomerase equivalent to the human immunophilin FKBP12, which also binds rapamycin. Therefore, we were interested in the possible interactions between rapamycin and echinocandins, particularly caspofungin.

Accordingly, this study represents the first demonstration of a synergistic relationship between rapamycin and caspofungin in the inhibition of the growth of *C. albicans* and *C. lusitaniae* strains sensitive and resistant to caspofungin. A positive interaction between these two molecules was reported about 20 years ago in *Aspergillus* species based on the agar disk diffusion method (23) but it was not subsequently confirmed via the checkerboard dilution method (24,25). This was likely because analysis of the results of a checkerboard test using the FICI is not suitable depending on the molecules tested, particularly when the MICs for one or both molecules involved in the association are low, which is the case here for rapamycin and caspofungin in *C. albicans* with MICs of 0.5 mg/L and 0.25 mg/L, respectively. In the present study, surface response analysis according to the Bliss model made it possible to detect the synergy for the two strains of *C. albicans* and to confirm the effect for the two strains of *C. lusitaniae*. This method seems to have greater discriminatory power because it is independent of the end point unlike the FICI method.

Synergy was observable not only for strains sensitive to caspofungin but also for resistant strains. For this demonstration, we used a clinical strain of *C. albicans* carrying one of the most common polymorphisms responsible for resistance to echinocandins in one allele of the *FKS1* gene, a nonsynonymous mutation in the HS1 region resulting in the S645P glucan synthase variant (26). This mutation is responsible for a decrease in binding affinity of the glucan synthase target to the inhibitor (27). We found that a rapamycin concentration as low as 0.125 mg/L was sufficient to restore caspofungin sensitivity. To confirm this observation, we introduced the equivalent S638P mutation in the Fks1 protein of a laboratory yeast model, *C. lusitaniae* (14), and demonstrated that rapamycin also restored sensitivity to caspofungin in an isogenic background. This indicates that the decrease in caspofungin binding affinity to glucan synthase was probably not the only mechanism involved in the resistance of strains with the S645P or equivalent substitution, and that there may be other mechanisms depending on the action of rapamycin.

From a molecular viewpoint, the synergy between rapamycin and caspofungin suggests that the two underlying signalling pathways, the TOR pathway for rapamycin (28,29) and the PKC pathway for caspofungin (30), are interconnected and share common effectors. In *Saccharomyces cerevisiae*, rapamycin activates the cell wall integrity (CWI) salvage pathway by phosphorylation of Slt2, the orthologous protein of Mkc1 in *C. albicans* (11). This observation strongly suggests crosstalk between the TOR and CWI pathways. Other work in *S. cerevisiae* has established a link between parietal stress induced by caffeine and Stl2 phosphorylation in a TOR-dependent manner (31). Other observations indicated that small G proteins from the RAS and Rho families may be good candidates at the intersection of these signalling pathways. In *C. albicans*, the Rhb1 protein belonging to the RAS superfamily could be involved in the activation of TOR because a homozygous mutant, *rhb1Δ/Δ*, showed increased sensitivity to rapamycin (32). In a more direct but also more complex way, it has been shown in *S. cerevisiae* that the Rho1 protein is involved in both negative regulation of the TORC1 complex and also that it could itself be activated by the TORC1 complex, depending on the environmental stresses to which yeasts are exposed (33). The information obtained in *S. cerevisiae* may not be directly transferrable to *C. albicans* because this species has only one copy of Tor, unlike *S. cerevisiae*, which has two Tor kinases (34). Nevertheless, Rho1 is located upstream of the PKC pathway. Reduced expression of *RHO1* in *C. albicans* (haploinsufficiency) results in increased susceptibility to caspofungin and calcofluor white (35,36). Rho1 has the dual function of regulating the glucan synthase protein complex and also of activating the CWI pathway under conditions of parietal stress, such as exposure to echinocandins (37,38).

The finding of synergistic interactions between caspofungin and rapamycin represents additional evidence for the involvement of the TOR pathway in the cellular response to parietal stress due to inhibition of glucan synthase by echinocandins.

## Transparency declarations

The authors have no conflicts of interest to declare.

## Author contributions

M. L., I. A. and T. N. designed the experiments. M. L. performed the experiments. M. L., I. A., N. B. and T. N. analysed the results. M. L., I. A. and T. N. wrote the manuscript.

## References

1 Brown GD, Denning DW, Gow NAR et al. Hidden killers: human fungal infections. Sci Transl Med. 2012; 4: 165rv13.

2 Pappas PG, Kauffman CA, Andes DR, et al. Clinical practice guideline for the management of candidiasis: 2016 update by the Infectious Diseases Society of America. Clin Infect Dis Off Publ Infect Dis Soc Am. 2016; 62: e1–50.

3 Denning DW. Echinocandin antifungal drugs. Lancet Lond Engl. 2003; 362: 1142–51.

4 Perlin DS. Mechanisms of echinocandin antifungal drug resistance. Ann N Y Acad Sci. 2015; 1354: 1–11.

5 Arendrup MC. Candida and candidaemia. Susceptibility and epidemiology. Dan Med J. 2013; 60: B4698.

6 Lackner M, Tscherner M, Schaller M et al. Positions and numbers of FKS mutations in Candida albicans selectively influence in vitro and in vivo susceptibilities to echinocandin treatment. Antimicrob Agents Chemother. 2014; 58: 3626–35.

7 Singh SD, Robbins N, Zaas AK et al. Hsp90 governs echinocandin resistance in the pathogenic yeast Candida albicans via calcineurin. PLoS Pathog 2009; 5: e1000532.

8 Zhang J, Silao FGS, Bigol UG et al. Calcineurin is required for pseudohyphal growth, virulence, and drug resistance in Candida lusitaniae. PLoS ONE 2012; 7: e44192.

9 Vézina C, Kudelski A, Sehgal SN. Rapamycin (AY-22,989), a new antifungal antibiotic. I. Taxonomy of the producing streptomycete and isolation of the active principle. J Antibiot. 1975; 28: 721–6.

10 Eng CP, Sehgal SN, Vézina C. Activity of rapamycin (AY-22,989) against transplanted tumors. J Antibiot. 1984; 37: 1231–7.

11 Torres J, Di Como CJ, Herrero E et al. Regulation of the cell integrity pathway by rapamycin-sensitive TOR function in budding yeast. J Biol Chem. 2002; 277: 43495–504.

12 Eltschinger S, Loewith R. TOR complexes and the maintenance of cellular homeostasis. Trends Cell Biol. 2016; 26: 148–159.

13 Tong Y, Zhang J, Wang L et al. Hyper-synergistic antifungal activity of rapamycin and peptide-like compounds against Candida albicans orthogonally via Tor1 kinase. ACS Infect Dis. 2021; 7: 2826–35.

14 Accoceberry I, Couzigou C, Fitton-Ouhabi V et al. Challenging SNP impact on caspofungin resistance by full-length FKS1 allele replacement in Candida lusitaniae. J Antimicrob Chemother. 2019; 74: 618–24.

15 Durrens P, Klopp C, Biteau N et al. Genome sequence of the yeast Clavispora lusitaniae type strain CBS 6936. Genome Announc. 2017; 5: e00724–17.

16 Clinical and Laboratory Standards Institute. Reference Method for Broth Dilution Antifungal Susceptibility Testing of Yeasts—Third Edition: Approved Standard M27-A3. CLSI, Wayne, PA, USA, 2008.

17 Odds FC. Synergy, antagonism, and what the chequerboard puts between them. J Antimicrob Chemother. 2003; 52: 1.

18 Di Veroli GY, Fornari C, Wang D et al. Combenefit: an interactive platform for the analysis and visualization of drug combinations. Bioinforma Oxf Engl. 2016; 32: 2866–8.

19 Bidaud AL, Djenontin E, Botterel F et al. Colistin interacts synergistically with echinocandins against Candida auris. Int J Antimicrob Agents. 2020; 55: 105901.

20 Denardi LB, Mario DAN, Loreto ÉS et al. Synergistic effects of tacrolimus and azole antifungal compounds in fluconazole-susceptible and fluconazole-resistant Candida glabrata isolates. Braz J Microbiol. 2015; 46: 125–9.

21 Cordeiro R de A, Macedo R de B, Teixeira CEC et al. The calcineurin inhibitor cyclosporin A exhibits synergism with antifungals against Candida parapsilosis species complex. J Med Microbiol. 2014; 63: 936–44.

22 Juvvadi PR, Lee SC, Heitman J et al. Calcineurin in fungal virulence and drug resistance: Prospects for harnessing targeted inhibition of calcineurin for an antifungal therapeutic approach. Virulence. 2016; 8: 186–97.

23 Kontoyiannis DP, Lewis RE, Osherov N et al. Combination of caspofungin with inhibitors of the calcineurin pathway attenuates growth in vitro in Aspergillus species. J Antimicrob Chemother. 2003; 51: 313–6.

24 Steinbach WJ, Schell WA, Blankenship JR et al. In vitro interactions between antifungals and immunosuppressants against Aspergillus fumigatus. Antimicrob Agents Chemother. 2004; 48: 1664–9.

25 Steinbach WJ, Singh N, Miller JL et al. In vitro interactions between antifungals and immunosuppressants against Aspergillus fumigatus isolates from transplant and nontransplant patients. Antimicrob Agents Chemother. 2004; 48: 4922–5.

26 Perlin DS. Current perspectives on echinocandin class drugs. Future Microbiol. 2011; 6: 441–57.

27 Perlin DS. Echinocandin Resistance in Candida. Clin Infect Dis Off Publ Infect Dis Soc Am. 2015; 61: S612–617.

28 Bastidas RJ, Heitman J, Cardenas ME. The protein kinase Tor1 regulates adhesin gene expression in Candida albicans. PLoS Pathog. 2009; 5: e1000294.

29 Rohde JR, Bastidas R, Puria R et al. Nutritional control via Tor signaling in Saccharomyces cerevisiae. Curr Opin Microbiol. 2008; 11: 153–60.

30 LaFayette SL, Collins C, Zaas AK et al. PKC signaling regulates drug resistance of the fungal pathogen Candida albicans via circuitry comprised of Mkc1, Calcineurin, and Hsp90. PLOS Pathog. 2010; 6: e1001069.

31 Kuranda K, Leberre V, Sokol S et al. Investigating the caffeine effects in the yeast Saccharomyces cerevisiae brings new insights into the connection between TOR, PKC and Ras/cAMP signalling pathways. Mol Microbiol. 2006; 61: 1147–66.

32 Tsao CC, Chen YT, Lan CY. A small G protein Rhb1 and a GTPase-activating protein Tsc2 involved in nitrogen starvation-induced morphogenesis and cell wall integrity of Candida albicans. Fungal Genet Biol. 2009; 46: 126–36.

33 Yan G, Lai Y, Jiang Y. The TOR complex 1 is a direct target of Rho1 GTPase. Mol Cell. 2012; 45: 743–53.

34 Qi W, Acosta-Zaldivar M, Flanagan PR et al. Stress- and metabolic responses of Candida albicans require Tor1 kinase N-terminal HEAT repeats. PLoS Pathog. 2022; 18: e1010089.

35 Edlind TD, Henry KW, Vermitsky JP et al. Promoter-dependent disruption of genes: simple, rapid, and specific PCR-based method with application to three different yeast. Curr Genet. 2005; 48: 117–25.

36 Corvest V, Bogliolo S, Follette P et al. Spatiotemporal regulation of Rho1 and Cdc42 activity during Candida albicans filamentous growth. Mol Microbiol. 2013; 89: 626–48.

37 Healey KR, Jimenez Ortigosa C, Shor E et al. Genetic drivers of multidrug resistance in Candida glabrata. Front Microbiol. 2016; 7: 1995.

38 Wiederhold NP, Kontoyiannis DP, Prince RA et al. Attenuation of the activity of caspofungin at high concentrations against Candida albicans: possible role of cell wall integrity and calcineurin pathways. Antimicrob Agents Chemother. 2005; 49: 5146–8.

